# *ATAD1* Overexpression Enhances Mitochondrial and Peroxisomal Function in Zellweger Syndrome Disorder Models

**DOI:** 10.1101/2025.10.14.682039

**Authors:** Diego Baronio, Tamara J. Stevenson, Courtney E. Demmitt-Rice, Esther C. Nuebel, Amanda M. Blackwell, Joshua L. Bonkowsky

**Affiliations:** Division of Pediatric Neurology, Department of Pediatrics, University of Utah School of Medicine, Salt Lake City, Utah; Department of Neurobiology, University of Utah School of Medicine, Salt Lake City, Utah; Department of Biochemistry, University of Utah School of Medicine, Salt Lake City, Utah; Roseman University of Health Sciences, College of Medicine, Henderson, Nevada; Center for Personalized Medicine, Primary Children’s Hospital, Salt Lake City, Utah

**Author notes:** Corresponding author: Josh Bonkowsky, Division of Pediatric Neurology, Department of Pediatrics, University of Utah 295 Chipeta Way, Salt Lake City, Utah, 84108. Equal contributors (Diego Baronio, Tamara J. Stevenson). Disclosures: JLB: has grants from NIH; clinical trials with Spur Therapeutics, Calico, and Ionis; consulting with Ionis; writing content for UpToDate; stock in Orchard Therapeutics; and royalties from BioFire and wFluidx. The University of Utah has filed a patent related to ATAD1, for which ECN and JLB are listed as co-inventors.

**Keywords:** Peroxisome, Mitochondria, Zellweger Syndrome, ATAD1, PEX

## Abstract

Zellweger Spectrum Disorders (ZSDs) are caused by mutations in any of the different peroxin (PEX) genes, which are essential for peroxisome biogenesis and function. Clinical features of ZSDs include seizures, leukodystrophy, renal and liver dysfunction, skeletal abnormalities, and they usually result in death during infancy or early childhood. There are no treatments for ZSDs, and their rarity, the large size of the PEX genes, and the numerous different genes, has impaired therapeutic development. We previously demonstrated that ATAD1, a mitochondrial protein quality control chaperone, could correct both mitochondrial and peroxisomal phenotypes in PEX3 patient fibroblasts. In this study, we investigated whether overexpressing ATAD1 could provide similar benefits in PEX1 and PEX6 patient cell lines, which account for over 70% of ZSD cases. We used established PEX6^-/-^ HEK293 cells, patient-derived fibroblasts with pathogenic PEX1 mutations, and newly created zebrafish mutants. Lipidomic profiling of the cell lines demonstrated widespread dysregulation, including accumulation of lysophosphatidylcholines with very-long-chain fatty acids, depletion of plasmalogens and cholesteryl esters containing polyunsaturated fatty acids, and a decrease in cardiolipins. Overexpressing ATAD1 partly corrected these imbalances, including normalizing VLCFA metabolism in PEX1 fibroblasts and restoring plasmalogens and cardiolipins in PEX6-deficient cells. Mitochondrial function analysis (Seahorse XF) showed that ATAD1 increased basal and ATP-linked respiration in both PEX1- and PEX6-deficient cells, sometimes surpassing the effects of PEX gene re-expression. ATAD1 increased peroxisome numbers in both PEX6 and PEX1 cells. Zebrafish Pex1 mutants exhibited impaired maximal respiration despite normal basal activity, confirming mitochondrial vulnerability in vivo. These findings further confirm a role for ATAD1 as a modifier that improves lipid metabolism, mitochondrial function, and peroxisome abundance that could function across multiple ZSDs.

## INTRODUCTION

Peroxisomal biogenesis disorders (PBDs) are rare autosomal recessive diseases caused by mutations in PEX genes, which encode peroxins—proteins essential for forming and maintaining peroxisomes^1^. The most severe PBDs are classified as Zellweger Spectrum Disorders (ZSDs), which feature seizures, leukodystrophy, hepatomegaly, kidney cysts, skeletal abnormalities, sensory deficits, and early childhood death^1,2^. Diagnosis relies on clinical signs and identifying biallelic pathogenic variants in one of the PEX genes^2^.

Peroxisomes are single-membrane organelles essential for metabolic processes, including fatty acid α- and β-oxidation, bile acid synthesis, glyoxylate detoxification, and plasmalogen biosynthesis^3^. They contain oxidative enzymes, such as catalase, which decomposes hydrogen peroxide^4^. Since peroxisomes are the sole site of plasmalogen production, a class of phospholipid ethers required for myelin synthesis, their malfunction primarily affects the central nervous system, leading to leukodystrophy and neurodegeneration^3^. A key biochemical marker of ZSD is the accumulation of very-long-chain fatty acids (VLCFAs), particularly C24:0 and C26:0, resulting from defective β-oxidation^5^.

Besides peroxisomal defects, mitochondrial issues are a hallmark of ZSD, first observed in liver tissue^6^. These problems arise not only from disrupted peroxisomal metabolism but also from mislocalized peroxins that insert into mitochondria, disturbing cristae structure and function. Recently we demonstrated that overexpressing ATAD1, a mitochondrial AAA-ATPase involved in membrane protein quality control, can partially restore mitochondrial shape and function in PEX3-deficient fibroblasts, suggesting that it may be a therapeutic target^7^.

ZSDs occur in about 1 in 50,000 births, and current treatments are purely palliative^1,2^. Among animal models, the PEX1 G844D mouse—mimicking the common human G843D mutation—demonstrates key disease features, such as hepatomegaly, retinal degeneration, mitochondrial malfunction, and early death, making it a valuable model for preclinical research^8,9^.

While gene replacement is a potentially attractive approach for ZSDs, most PEX genes are too large for packaging in adeno-associated virus (AAV) vectors^10,11^. Furthermore, therapies would need to be specifically developed and tested for each of the 14 different PEX genes. Conversely, ATAD1 offers two main advantages: its small coding size (∼1 kb) allows efficient AAV delivery, and its downstream role could provide functional rescue across many PEX mutations. As a result, ATAD1-based therapy might be a more universal and scalable strategy for ZSDs.

Due to the clinical and genetic variability of ZSDs, our goal was to evaluate the therapeutic potential of ATAD1 overexpression in additional models with various PEX mutations. Specifically, we analyzed metabolic and mitochondrial outcomes in a patient-derived fibroblast with a PEX1 mutation and a CRISPR-engineered HEK293 cell line with a PEX6 mutation^12^—the two most frequently affected genes in ZSD. We also tested whether ATAD1 overexpression could improve mitochondrial function and reduce VLCFA accumulation. Finally, we extended our study to pex1 and pex6 zebrafish mutants to evaluate behavioral traits and mitochondrial activity *in vivo*, providing further insights into ZSD pathology and potential treatments.

## RESULTS

### Generation and Validation of PEX-deficient models

To examine the effects of ATAD1 in various Zellweger Spectrum Disorder (ZSD) models, we used two human cell lines: HEK293 cells lacking PEX6, created through CRISPR/Cas9 editing^12^, and fibroblasts from a patient with compound heterozygous pathogenic PEX1 mutations. The PEX6 mutant line has a single-nucleotide deletion (c.544delG) that causes a frameshift after valine 181, resulting in a premature stop at residue 204 and loss of the N2, D1, and D2 functional domains. In the PEX1-deficient fibroblasts, the first mutation is a single-nucleotide duplication (c.2097dupT) that introduces a frameshift after phenylalanine 699 and a premature stop at residue 740, disrupting the C-terminal part of the D1 ATPase domain and removing the D2 domain. The second mutation (c.2528G>A) causes a missense change from glycine 843 to aspartic acid (G843D) within the D2 ATPase domain. This well-characterized hypomorphic allele impairs PEX1 folding and ATPase activity but retains some function^13,14^.

The positions of the PEX1 I700fs (c.2097dupT), PEX1 G843D (c.2528G>A), and PEX6 c.544delG mutations are shown within the relevant gene structures (Figure 1A). We refer to these lines as PEX1^-/-^ and PEX6^-/-^ throughout the paper; mutations were confirmed by sequencing at the genomic locus and of the mRNA (data not shown). Alphafold structural predictions revealed that all mutations leading to early protein termination eliminated essential functional domains of PEX1 and PEX6, respectively (Figure 1B–D).

**Figure 1.**
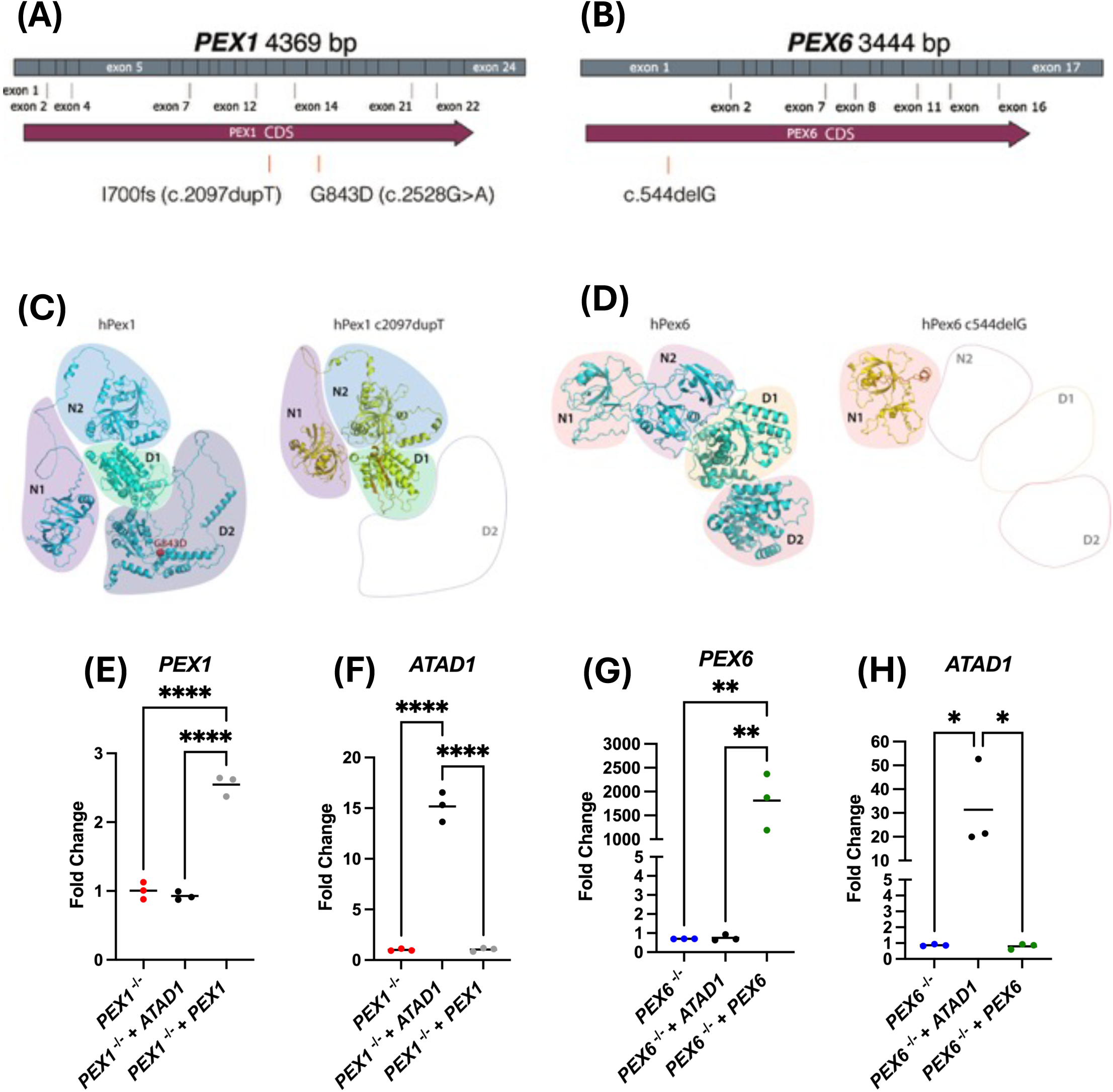
Generation and characterization of PEX1 and PEX6 mutant cell models. (A) Schematic representation of *PEX1* and *PEX6* gene variants. The *PEX1*-deficient fibroblasts harbor compound heterozygous mutations: c.2097dupT (I700fs), truncating the D2 domain, and c.2528G>A (G843D), a hypomorphic missense variant within the D2 ATPase domain. The *PEX6* mutant HEK293 line harbors a homozygous c.544delG frameshift mutation, resulting in premature termination and loss of the N2, D1, and D2 domains. (**B–D**) Alphafold structural models of wild-type and mutant PEX1 and PEX6 proteins showing disrupted AAA+ ATPase domains or altered folding caused by the respective mutations. (**E–H**) qRT-PCR analysis confirming overexpression of *PEX1*, *PEX6*, and *ATAD1* following transfection in mutant cell lines. Dot plots show fold-changes. Data is represented as mean ± SE. Statistical analysis used One-way ANOVA with Tukey’s multiple comparisons test. *P ≤ 0.05, **P ≤ 0.01, ****P ≤ 0.0001

Next, we confirmed the overexpression of rescue constructs in these models. Quantitative RT-PCR revealed that mutant cells had significantly increased levels of PEX1, PEX6, and ATAD1 transcripts (respectively) after transduction (Figure 1E–H). These results establish the genetically defined cell lines and the transduction approach for testing effects.

### ATAD1 Overexpression Restores Lipid Homeostasis in PEX1- and PEX6-Deficient Cells

Aberrant accumulation of peroxins on mitochondria in ZSDs results in pronounced defects in mitochondrial morphology, respiration, and lipid metabolism. Notably, these abnormalities can be ameliorated by the overexpression of ATAD1^7^. Recent studies have emphasized the intricate interplay between mitochondrial and peroxisomal metabolism, encompassing not only physical contact sites and very-long-chain fatty acid (VLCFA) β-oxidation, but also plasmalogen biosynthesis^15,16^. It has been proposed that perturbations in mitochondrial metabolism, structure, and respiratory function may compromise the synthesis of lipid species critical for mitochondrial integrity, such as cardiolipins, as well as phosphor-ether-lipids in the peroxisome. Given that phosphoether-lipids serve as essential precursors for myelin formation, we investigated whether overexpression of ATAD1 in our cellular models could modulate the abundance of these specific lipid classes.

To determine the effects of ATAD1 overexpression on lipid abnormalities observed in ZSDs, we conducted untargeted lipidomic analyses on PEX1^-/-^ fibroblasts and PEX6^-/-^ HEK293 cells. Global profiling revealed widespread changes across multiple lipid classes in both models, which were partially corrected by ATAD1 overexpression (Figure 2A and 2 B).

**Figure 2.**
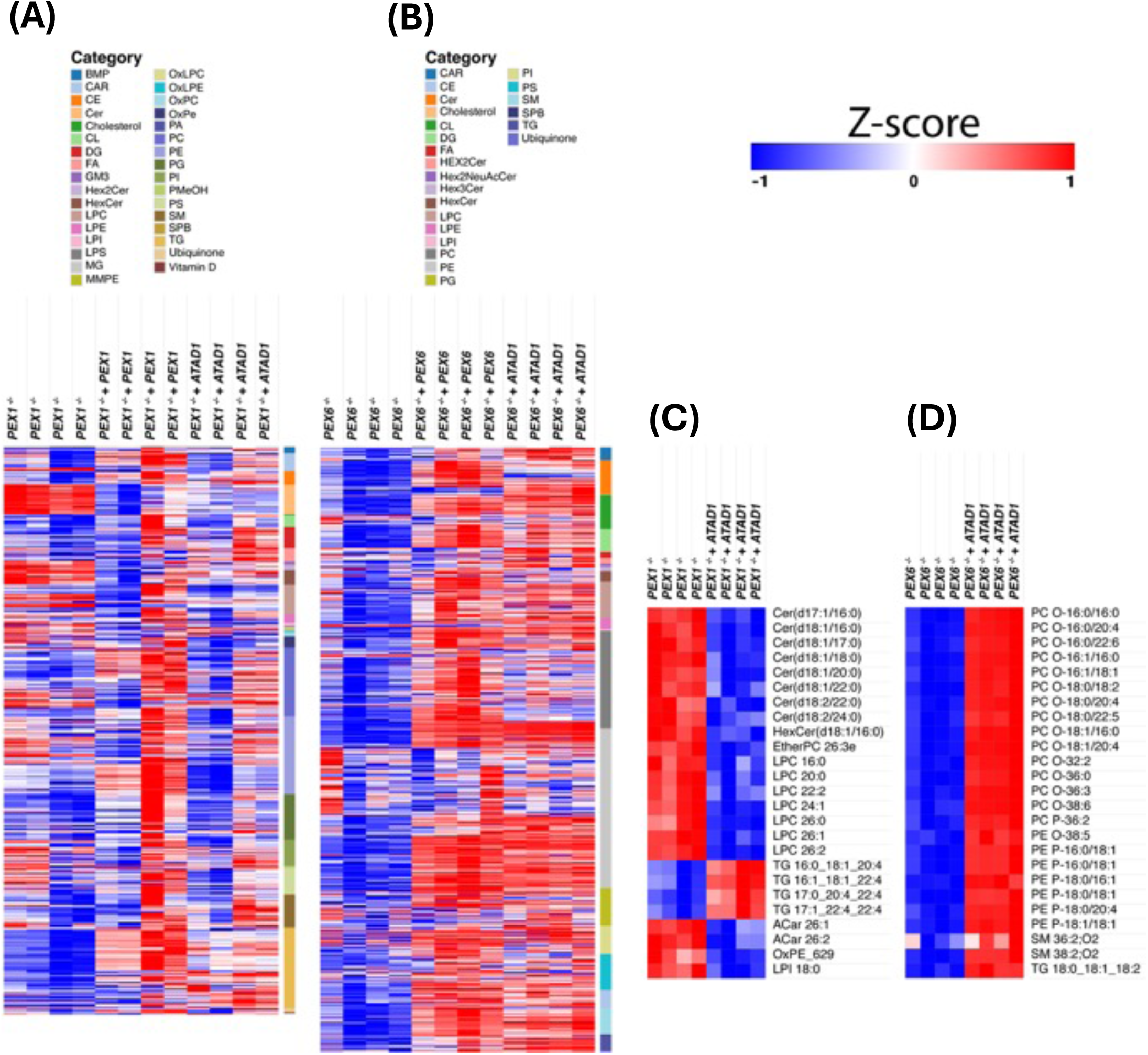
ATAD1 rescues lipidomic abnormalities in PEX1⁻/⁻ and PEX6⁻/⁻ cell models. (**A–B**) Heatmaps of untargeted lipidomic profiles in PEX1⁻/⁻ fibroblasts (**A**) and PEX6⁻/⁻ HEK293 cells (**B**), compared to their respective PEX1- and PEX6-rescued controls, with and without ATAD1 overexpression. Lipid species are grouped by class (color-coded by category on the right), and values are presented as Z-scores (blue = decreased, red = increased). Both gene rescue and ATAD1 partially normalize lipid alterations, with overlapping but distinct effects. (**C–D**) Top 25 lipid species most affected by ATAD1 in PEX1**⁻/⁻** (**C**) and PEX6**⁻/⁻** (**D**) cells. In PEX1⁻/⁻ cells, ATAD1 reduces very-long-chain LPCs and increases PUFA-TGs, suggesting improved β-oxidation. In PEX6⁻/⁻ cells, ATAD1 increases plasmalogens and sphingomyelins, indicating restored ether lipid synthesis and membrane organization.

To identify the lipid species most responsive to ATAD1, we analyzed the top 25 lipids with the most significant differential abundance (Figures 2C and 2D). In PEX1^-/-^ fibroblasts, levels of very-long-chain lysophosphatidylcholines (LPCs), which indicate impaired peroxisomal β-oxidation, were markedly elevated, while ATAD1 expression significantly reduced these levels. Conversely, polyunsaturated triacylglycerols (PUFA-TGs) levels increased with ATAD1, implying enhanced incorporation or availability of fatty acids from peroxisomes. In PEX6^-/-^ cells, ATAD1 led to increases in plasmalogens, suggesting a restoration of ether lipid synthesis, as well as elevated sphingomyelin species, indicating improved membrane organization and lipid trafficking. Overall, these findings demonstrate that ATAD1 overexpression rectifies key lipid dysregulation features in both PEX1^-/-^ and PEX6^-/-^ cells, although the relative amounts of specific lipid classes affected varied.

The different responses to ATAD1 overexpression in PEX1^-/-^ and PEX6^-/-^ cells may reflect model-specific factors as well as gene-specific factors. PEX1 modeling was performed in patient fibroblasts, whereas for PEX6 studies were in a HEK293 line. Although both PEX1 and PEX6 encode interacting AAA ATPases essential for peroxisomal matrix protein import, they are not interchangeable^17^. PEX1 has a more prominent role in unfolding substrates and recycling the import receptor PEX5, and its deficiency generally leads to more severe biochemical and clinical outcomes compared to PEX6 deficiency^18,19^. These inherent differences could influence how peroxisomal dysfunction disrupts lipid metabolism and how responsive the system is to secondary rescue by ATAD1.

Building on the broad lipid remodeling shown by the heatmaps, we then conducted detailed targeted lipidomic analyses to measure specific lipid classes altered in PEX6^-/-^ and PEX1^-/-^ cells and evaluate the effects of ATAD1 or PEX gene overexpression (Figure 3).

**Figure 3.**
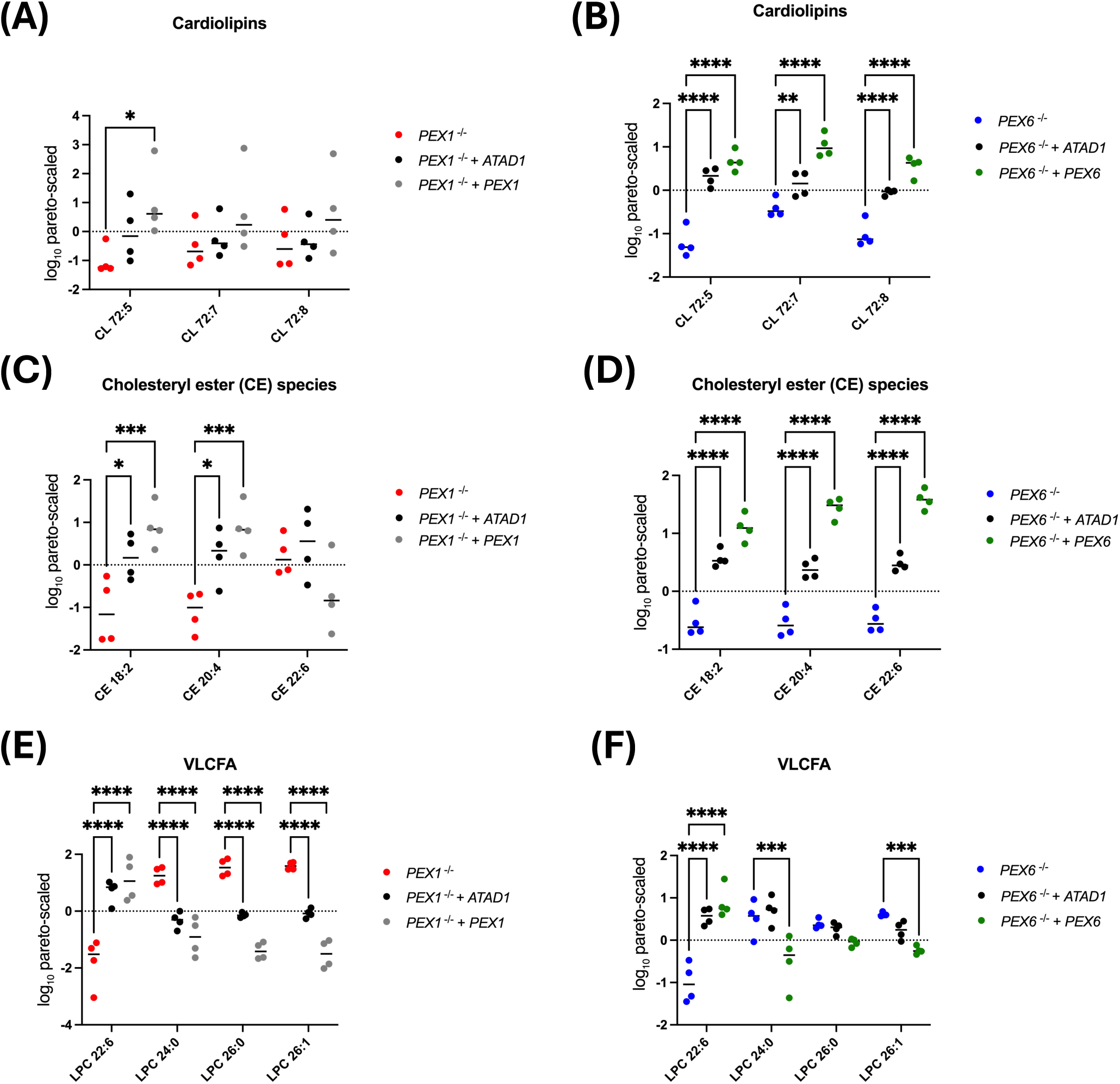
Targeted lipidomics reveals selective lipid rescue by ATAD1 or PEX gene expression. (**A–B**) Cardiolipins (CL 72:5, 72:7, 72:8) are decreased in PEX1⁻/⁻ and PEX6⁻/⁻ cells. PEX6 or ATAD1 fully restores these in PEX6⁻/⁻, while only CL 72:5 is recovered by PEX1 in PEX1⁻/⁻; ATAD1 shows no effect. (**C–D**) PUFA-rich cholesteryl esters (CE 18:2, 20:4, 22:6) are reduced in both models. CE 18:2 and 20:4 are rescued by PEX1 or ATAD1 in PEX1⁻/⁻; all three are restored in PEX6⁻/⁻ by PEX6 or ATAD1. (**E–F**) VLCFA-containing LPCs (24:0, 26:0, 26:1) accumulate in PEX1⁻/⁻ and are reduced by PEX1 or ATAD1. LPC 22:6 is partially restored. In PEX6⁻/⁻, ATAD1 improves LPC 22:6 but not VLCFA-LPCs. Dot plots show log-transformed, Pareto-scaled data as mean ± SE. Statistical analysis used One-way ANOVA with Tukey’s multiple comparisons test. *P ≤ 0.05, **P ≤ 0.01, ***P ≤ 0.001, ****P ≤ 0.0001

In PEX6^-/-^ cells, the levels of cardiolipins CL 72:5, CL 72:7, and CL 72:8 were notably reduced, indicating damage to mitochondrial inner membrane integrity. Restoring PEX6 expression or overexpressing ATAD1 fully recovered these three cardiolipin species (Figure 3B). These results reveal that ATAD1 can enhance cardiolipin remodeling but does not entirely substitute for PEX6’s function. In PEX1^-/-^ fibroblasts, the same cardiolipins were also decreased, but the pattern of rescue differed. Reintroducing PEX1 restored only CL 72:5, whereas ATAD1 overexpression did not recover any of the species (Figure 3A). This suggests that ATAD1’s capacity to compensate for cardiolipin metabolism is more effective in the PEX6^-/-^ context than in the PEX1^-/-^ one.

In PEX6^-/-^ cells, levels of cholesteryl esters (CEs) (Figure 3D) with polyunsaturated fatty acids (PUFAs)—notably CE 18:2, CE 20:4, and CE 22:6—were significantly decreased. These compounds are key indicators of lipid metabolic health: CE 18:2 (linoleic acid) and CE 20:4 (arachidonic acid) serve as precursors for bioactive lipids involved in inflammation, while CE 22:6, an ester of docosahexaenoic acid (DHA), acts as a circulating reservoir supporting membrane fluidity and neuroprotection by providing DHA for membrane phospholipids^20,21^. In PEX1^-/-^ cells (Figure 3C), levels of CE 18:2 and CE 20:4 were restored both by PEX1 and ATAD1, but CE 22:6 levels showed no significant differences across groups. Considered together, plasmalogens and an increase in CE lipid classes are interpreted as indicating active peroxisomal lipid metabolism, with improved flux towards both ether lipids and esterified cholesterol storage. Reintroducing PEX6 or expressing ATAD1 significantly increased the levels of these PUFA-rich CEs compared to the mutant, indicating improved peroxisomal function, PUFA metabolism, and cholesterol esterification.

Accumulation of VLCFA-containing LPCs is a hallmark of impaired peroxisomal β-oxidation and is a biochemical marker for ZSD^5^. Notably, VLCFAs, especially when esterified in lysophosphatidylcholine forms like LPC 24:0, 26:0, and 26:1, were significantly elevated in PEX1^-/-^ fibroblasts (Figure 3E). This increase reflects defective peroxisomal β-oxidation and abnormal fatty acid catabolism typical of ZSD. The buildup of VLCFA-rich LPCs signals a disruption in peroxisomal pathways responsible for VLCFA degradation. Conversely, LPC 22:6 (which contains docosahexaenoic acid) was notably reduced in PEX1^-/-^ cells, indicating a specific impairment in DHA metabolism—an essential factor for membrane fluidity and neuroprotection. Importantly, overexpressing ATAD1 or PEX1 in PEX1^-/-^ fibroblasts significantly lowered these elevated VLCFA-LPC levels and partially restored LPC 22:6 concentrations to control levels. In PEX6^-/-^ cells (Figure 3F), ATAD1 failed to reduce the levels of LPC 24:0, 26:0, and 26:1. However, the depletion of LPC 22:6 seen in mutant cells was attenuated by ATAD1 treatment. These findings suggest that ATAD1 plays a role in restoring peroxisomal fatty acid processing and lipid balance, offering potential for correcting lipid imbalances linked to ZSDs.

### ATAD1 Overexpression Restores Mitochondrial Function Assessed by Seahorse XF Analysis

To investigate how peroxisomal dysfunction impacts mitochondrial bioenergetics and whether this can be mitigated by gene rescue, we performed Seahorse XF assays on PEX1^-/-^ or PEX6^-/-^ cells (Figures 4A and 5A). These cells were compared to those that either re-expressed the respective PEX gene or overexpressed ATAD1.

**Figure 4.**
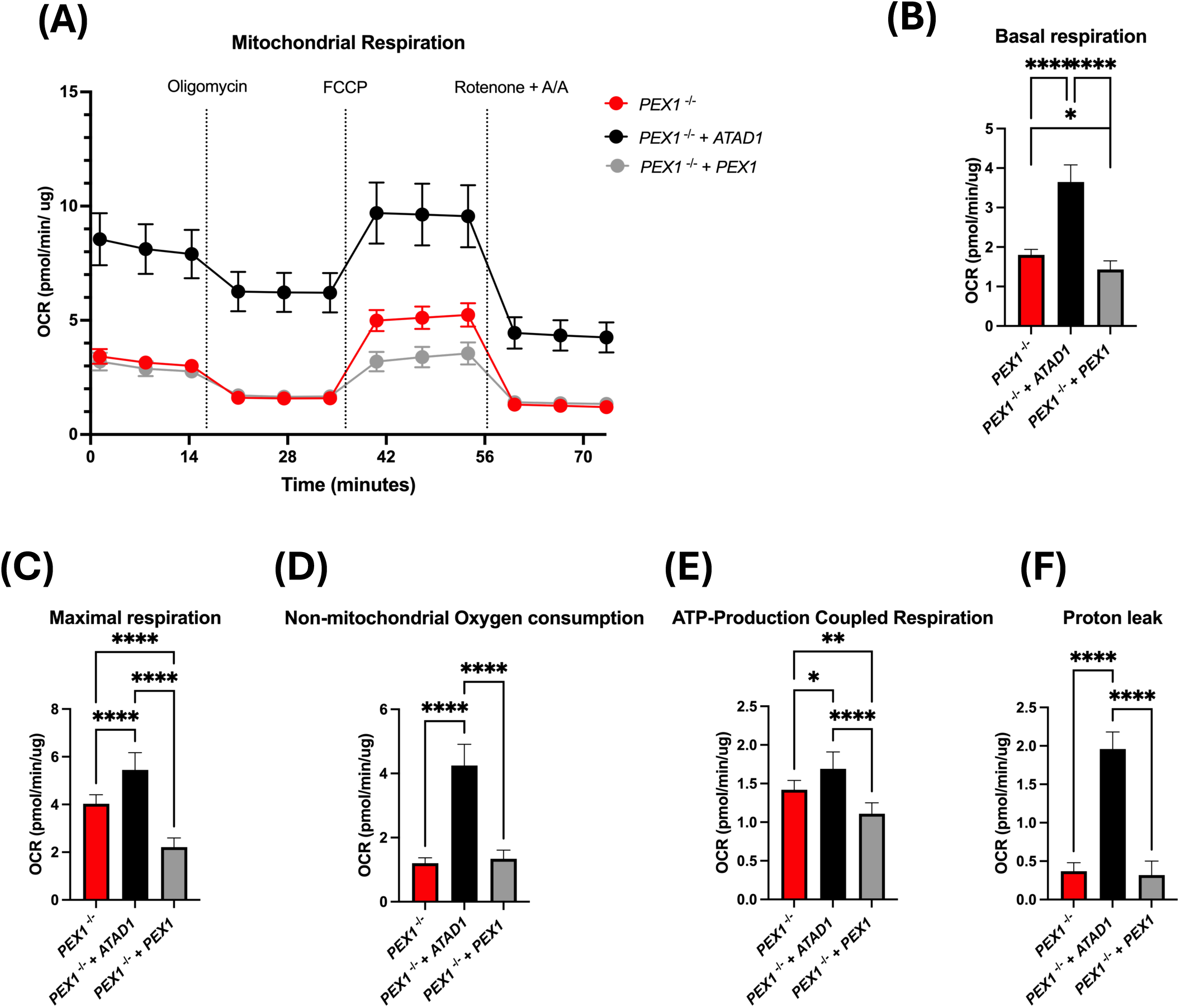
ATAD1 overexpression enhances mitochondrial respiration in PEX1-deficient fibroblasts. **(A)** Seahorse XF assay measuring OCR (pmol/min/ug) over time in PEX1-/- fibroblasts (red), PEX1-/- cells re-expressing PEX1 (grey), and PEX1-/- cells overexpressing ATAD1 (black). Key respiratory modulators were added as indicated: oligomycin (Oligo), FCCP, and rotenone/antimycin A (Rot/AA). (**B-E**) Quantification of mitochondrial respiration parameters derived from the Seahorse assay: (B) Basal respiration, (**C**) Maximal respiration, (**D**) Non-mitochondrial respiration, (**E**) ATP-coupled respiration, and (F) Proton leak. Statistical analysis used One-way ANOVA with Tukey’s multiple comparisons test. Data are mean ± SEM; n = 3 independent experiments. *p ≤ 0.05, **p ≤ 0.01, ****p ≤ 0.0001.

**Figure 5.**
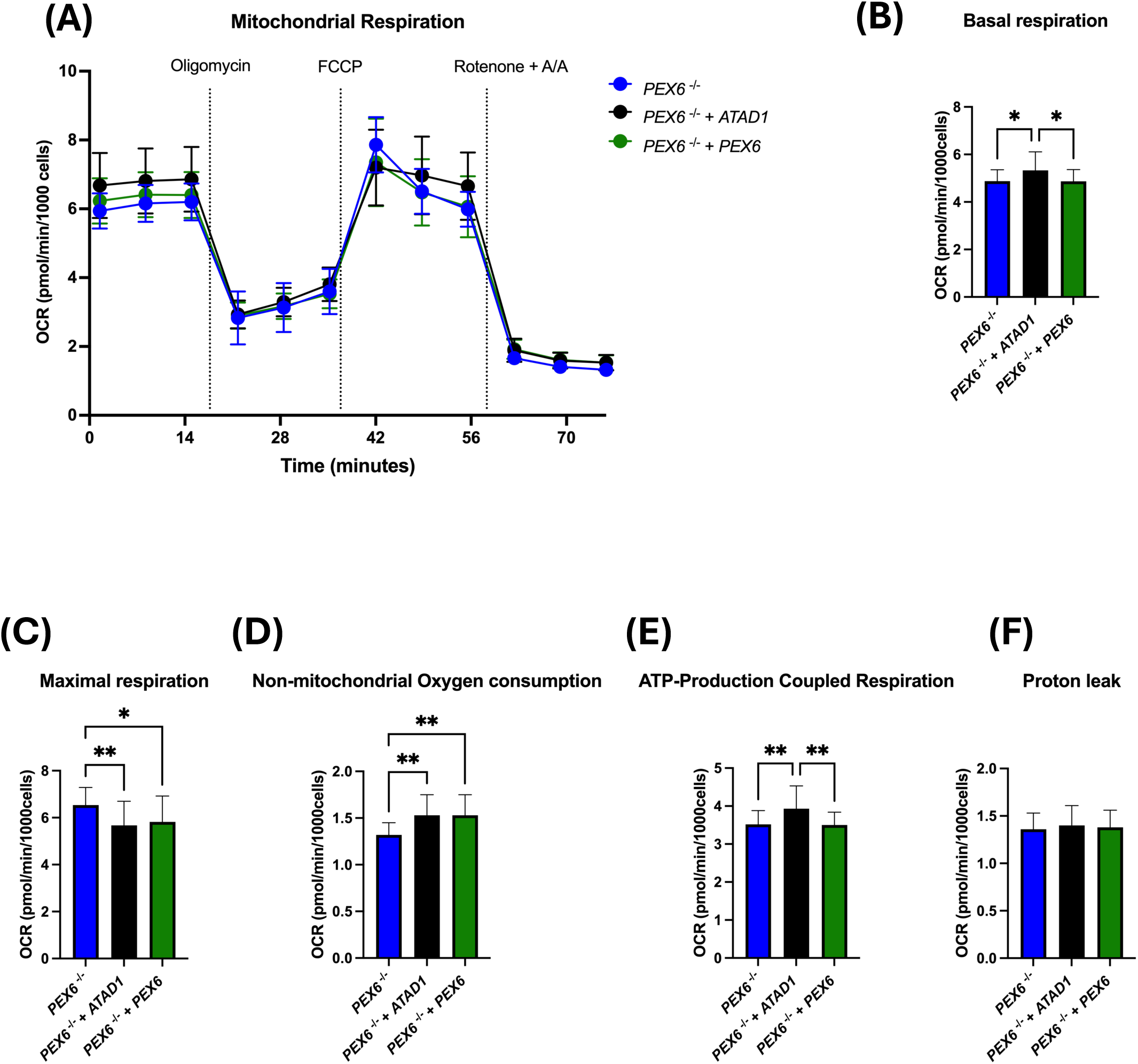
ATAD1 overexpression enhances mitochondrial respiration in PEX6-deficient HEK293 cells. **(A)** Seahorse XF assay measuring OCR (pmol/min/1000 cells) over time in PEX6-/- cells (blue), PEX6-/- cells re-expressing PEX6 (green), and PEX6-/- cells overexpressing ATAD1 (black). Key respiratory modulators were added as indicated: oligomycin (Oligo), FCCP, and rotenone/antimycin A (Rot/AA). (**B-E**) Quantification of mitochondrial respiration parameters derived from the Seahorse assay: (B) Basal respiration, (**C**) Maximal respiration, (**D**) Non-mitochondrial respiration, (**E**) ATP-coupled respiration, and (**F**) Proton leak. Statistical analysis used One-way ANOVA with Tukey’s multiple comparisons test. Data are mean ± SEM; n = 3 independent experiments. *p ≤ 0.05, **p ≤0.01.

In PEX1^-/-^ fibroblasts, overexpression of ATAD1 increased basal respiration compared to both PEX1^-/-^ cells and PEX1-rescued cells (Figure 4B). Conversely, reintroducing PEX1 worsened this parameter in the mutant cells. A similar pattern was observed with ATP-coupled respiration, where ATAD1 overexpression significantly improved function, whereas re-expression of PEX1 failed to restore it to a significant extent (Figure 4E).

Proton leak was significantly increased by ATAD1 overexpression, while PEX1 re-expression had no effect (Figure 4F). This increase in proton leak may indicate enhanced mitochondrial activity but also suggests greater proton conductance, possibly due to higher respiratory chain activity or slight mitochondrial uncoupling.

Notably, maximal respiration was reduced by PEX1 re-expression (Figure 4C). In contrast, ATAD1 overexpression caused a significant increase in maximal respiration compared to both PEX1^-/-^ and PEX1-rescued cells. Lastly, non-mitochondrial oxygen consumption remained unaffected by PEX1 overexpression. However, it significantly increased in cells overexpressing ATAD1, indicating the activation of alternative oxygen-consuming pathways, possibly related to stress responses or redox shifts (Figure 4D).

In the context of PEX1 deficiency, ATAD1 overexpression not only restores basal and ATP-linked respiration but also enhances maximal capacity, increases proton leak, and raises non-mitochondrial oxygen consumption. These findings support a role for ATAD1 as a potent enhancer of mitochondrial bioenergetics, potentially by promoting mitochondrial protein quality control or dynamics in cells where peroxisomal function is impaired.

In PEX6^-/-^ cells, basal respiration (Figure 5A) was significantly increased by ATAD1 overexpression compared to both untreated mutants and PEX6-rescued cells. However, re-expression of PEX6 alone did not improve basal respiration, indicating that ATAD1—but not PEX6—can partially restore this key aspect of mitochondrial function.

A similar pattern was seen in ATP-linked respiration (Figure 5E), where ATAD1 significantly enhanced ATP production compared to both PEX6^-/-^ and PEX6^-/-^ + PEX6, reinforcing ATAD1’s role in increasing mitochondrial output. Meanwhile, proton leak (Figure 5F) stayed consistent across all groups, suggesting no notable alterations in mitochondrial membrane integrity or uncoupling.

Interestingly, the highest maximal respiration (Figure 5B) was observed in untreated PEX6^-/-^ cells and was notably diminished by both PEX6 re-expression and ATAD1 overexpression. These interventions seem to increase basal and ATP-linked respiration but may compromise the mitochondria’s ability to adapt to higher energetic demands, possibly due to altered mitochondrial dynamics. This indicates a shift toward more sustained yet less adaptable mitochondrial activity. Furthermore, non-mitochondrial oxygen consumption (Figure 3D) was significantly elevated in both rescue conditions—PEX6 and ATAD1—compared to mutants, with no significant difference between them. This increase might reflect the activation of alternative oxygen-consuming pathways, potentially as a compensatory mechanism in response to cellular stress or redox imbalance.

These findings imply that re-expressing PEX6 does not fully restore mitochondrial function in knockout cells. Nonetheless, ATAD1 provides a partial recovery by improving both basal and ATP-linked respiration. The decrease in respiratory reserve and increase in non-mitochondrial oxygen consumption suggest a metabolic shift, potentially in response to cellular stress or adaptation.

### ATAD1 Overexpression and PEX6 Re-expression Increase Peroxisome Numbers in PEX6^-/-^ Cells

To assess whether ATAD1 overexpression or PEX6 re-expression can restore peroxisome formation in PEX6^-/-^ cells, we performed immunofluorescence microscopy on our knockout and complement cell lines, staining for peroxisomal PMP70 (Figure 6A). Overexpressing ATAD1 in PEX6^-/-^ cells significantly increased the PMP70-positive area per cell. Reintroducing PEX6 led to the production of a similar phenotype, bringing the counts to levels like those in wild-type cells (Figure 6B). Measurement of the PMP70-stained area as a percentage of the cell area confirmed that the difference between groups persisted, indicating that peroxisome abundance was independent of cell volume (Figure 6C).

**Figure 6.**
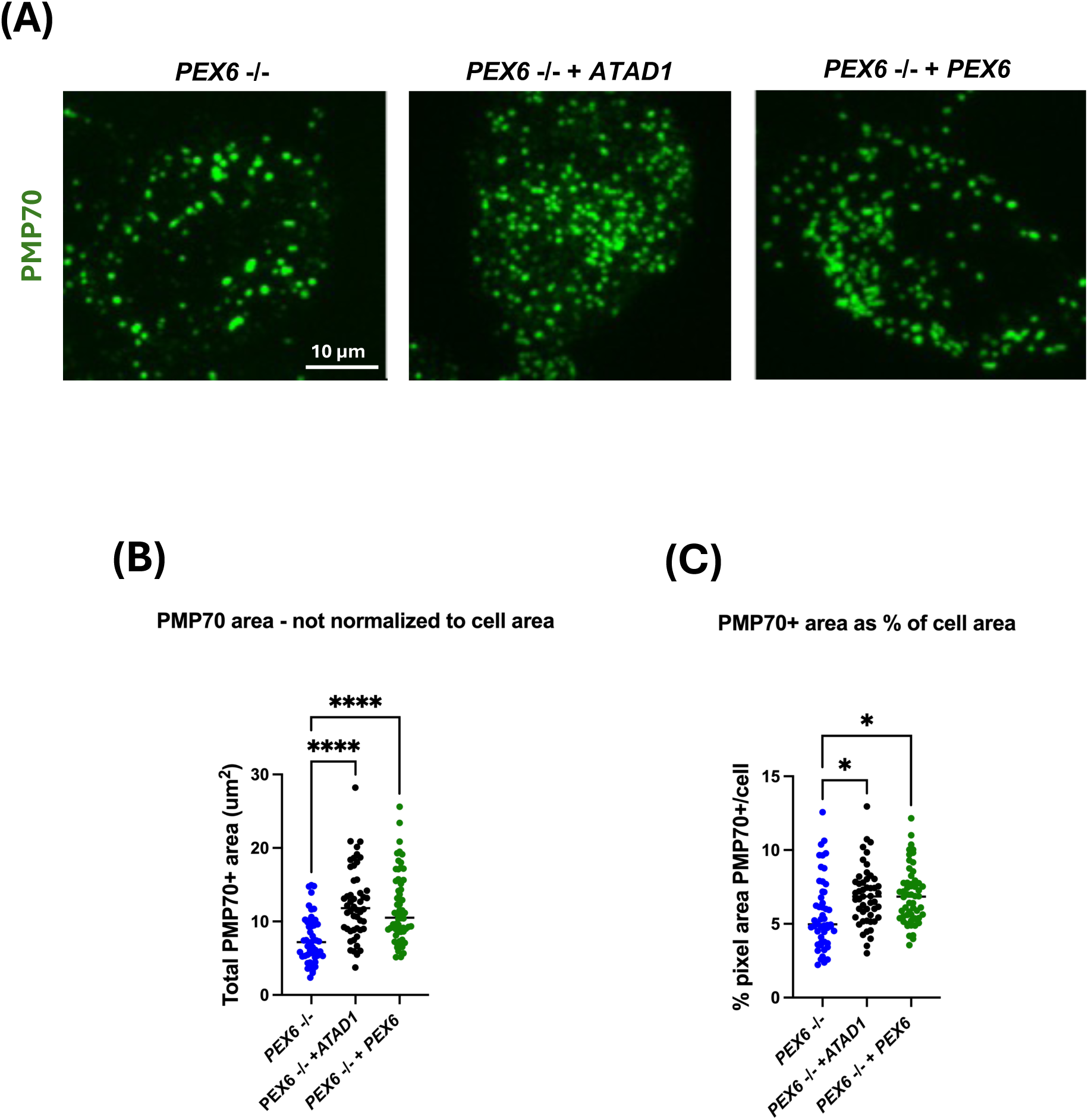
ATAD1 overexpression increases the number of peroxisomes in PEX6-deficient cells. **(A)** Representative immunofluorescence images of HEK293 PEX6-/- cells, PEX6-/- cells overexpressing ATAD1, and PEX6-/- cells re-expressing PEX6, stained for the peroxisomal membrane protein PMP70 (green). Images were captured using a Nikon Spinning Disk Confocal microscope. (Scale bar 10 μm). (**B**) The PMP70-positive area shows a significant increase upon ATAD1 overexpression and a further increase after PEX6 re-expression. (**C**) The PMP70-positive area, expressed as a percentage of total cell area, confirms that differences in peroxisome abundance are independent of cell size.

### Generation of zebrafish mutant lines and assessment of locomotor behavior and mitochondrial function

To study zebrafish models of peroxisomal dysfunction, we characterized mutant lines for Pex1 and Pex6. Using CRISPR/Cas9, we introduced a 44-base pair deletion in exon 5 of the *Pex1* (Figure 7A) gene (c.802del44), causing a frameshift at proline-267 that leads to an early termination after only 296 residues and eliminating the Pex1 N2, D1, and D2 functional domains (Figure 7B), as demonstrated by AlphaFold. A *Pex6* mutant line carrying a nucleotide substitution in exon 16 (c.2786C>A; genomic G>T), introducing a premature stop codon after histidine-928 causing premature termination of the normally 1077–amino acid Pex6 protein (Figure 7D), was obtained from the Zebrafish International Resource Center (ZIRC) and confirmed by sequencing at the gene locus as well as by sequencing of the cDNA. The Pex1 and Pex6 lines were crossed to produce double-heterozygous fish, which were bred to generate wild-type (WT; siblings of the double-mutants), heterozygous, and double-mutant larvae for further study. To evaluate the impact of Pex1 and/or Pex6 loss on early zebrafish behavior, we measured locomotor activity in 6-day post-fertilization (dpf) larvae (Figure 7B). Total distance traveled over 30 minutes was tracked using an automated system. No significant differences in total distance were found comparing wild-type, Pex1 homozygotes, Pex6 homozygotes, and double-mutant larvae. These findings suggest that, under normal conditions, disrupting Pex1 and Pex6 does not affect spontaneous locomotion at this stage of development.

**Figure 7.**
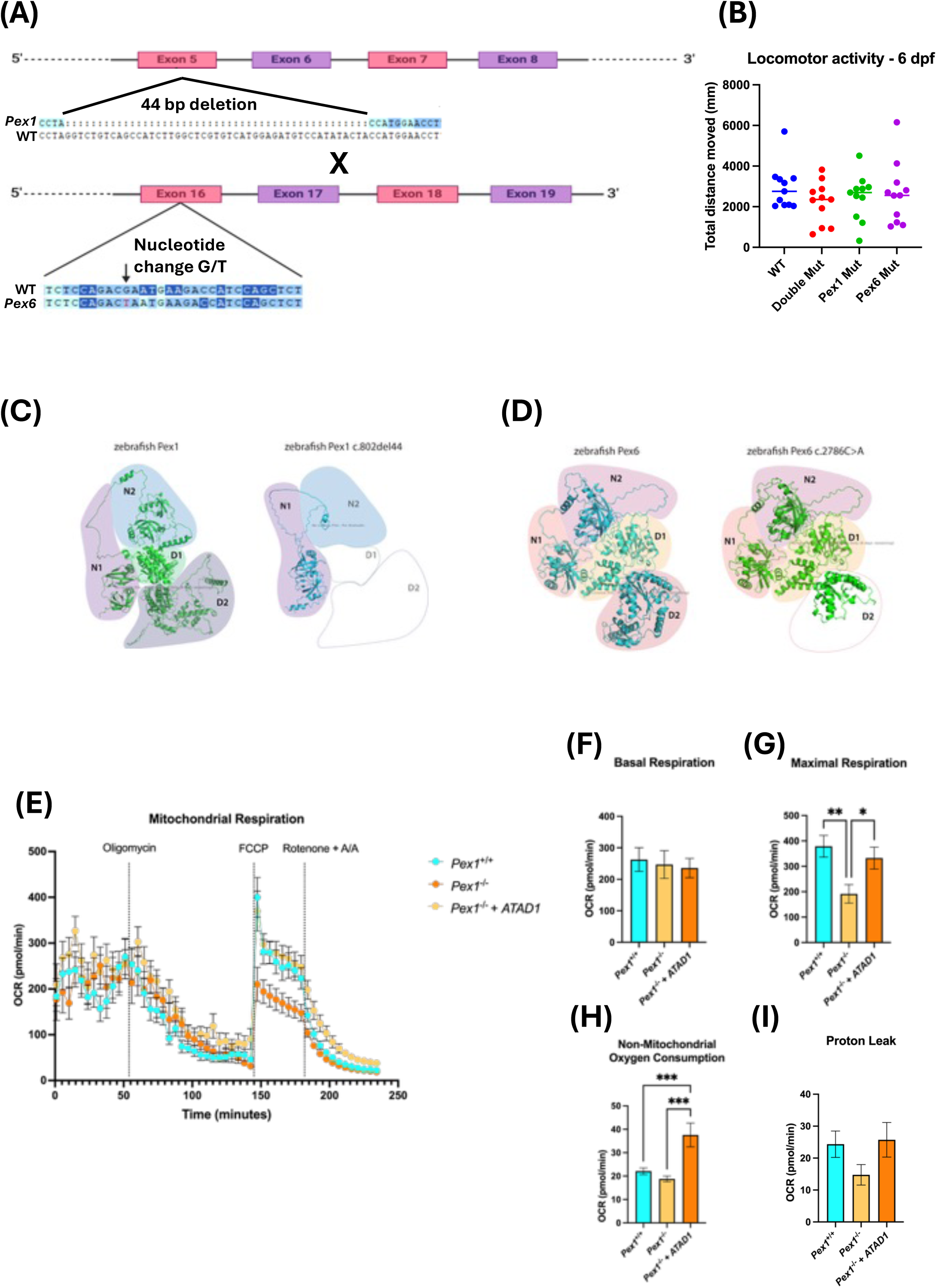
Generation and functional characterization of zebrafish *pex1* and *pex6* mutants. (A) Schematic representation of *pex1* (c.802del44) and *pex6* (c.2786C>A; genomic G>T) mutations introduced by CRISPR/Cas9 or obtained from ZIRC. (**B**) Locomotor activity of 6 dpf larvae from WT, *Pex1⁻/⁻*, *Pex6⁻/⁻*, and double-mutant lines; total distance traveled over 30 min showed no significant differences (n = 11 per group). (**C–D**) AlphaFold-predicted structures of WT and mutant Pex1 and Pex6 proteins. The *Pex1* mutation causes a frameshift at Pro267, resulting in the loss of the N2, D1, and D2 ATPase domains. In contrast, the *Pex6* mutation introduces a premature stop codon after His928, thereby truncating the normally 1077–amino acid protein. (**E**) Seahorse XF analysis of mitochondrial oxygen consumption rate (OCR) in WT, *Pex1⁻/⁻*, and *Pex1⁻/⁻ + ATAD1* zebrafish. (F–H) Quantification of basal, maximal, and non-mitochondrial respiration. *pex1⁻/⁻* mutants exhibited reduced maximal respiration, which was restored by ATAD1 expression. Statistical analysis used One-way ANOVA with Tukey’s multiple comparisons test. Data are mean ± SEM. *p ≤ 0.05, **p ≤0.01, ***P ≤ 0.001.

Mitochondrial respiration was evaluated in WT and Pex1^-/-^ mutant zebrafish using Seahorse XF analysis (Figure 7E). The basal respiration (Figure 7F) did not significantly differ among the three groups, indicating that resting mitochondrial activity is preserved in the Pex1⁻/⁻ mutants. Upon oligomycin injection, ATP-linked respiration decreased across all groups, consistent with inhibition of ATP synthase activity (Figure 7E). No significant differences were detected between WT, Pex1⁻/⁻, and Pex1⁻/⁻ + ATAD1 conditions at this stage. Following FCCP treatment, which uncouples oxidative phosphorylation to reveal maximal respiratory capacity, WT zebrafish showed a marked increase in OCR, whereas Pex1⁻/⁻ mutants exhibited a blunted response. Expression of ATAD1 in the mutant background restored this response to WT levels, with maximal respiration significantly increased compared to Pex1⁻/⁻ alone (Figure 7G). After rotenone and antimycin A treatment, OCR values dropped to levels corresponding to non-mitochondrial respiration (Figure 7H). These values were significantly higher in Pex1⁻/⁻ + ATAD1 zebrafish than in Pex1⁻/⁻ mutants, suggesting enhanced non-mitochondrial oxygen-consuming activity upon ATAD1 expression. Proton leak remained comparable across all groups, with only a nonsignificant trend toward reduction in the Pex1⁻/⁻ mutants (Figure 7I). Together, these results demonstrate that while Pex1 deficiency impairs mitochondrial reserve capacity, ATAD1 expression effectively restores mitochondrial respiratory function, normalizing both maximal and non-mitochondrial respiration to levels comparable to WT zebrafish.

## DISCUSSION

We found that overexpression of ATAD1, a mitochondrial chaperone protein, rescued key Zellweger Syndrome Disorder (ZSD)–associated biomarkers, including reduction of elevated very long-chain fatty acids (VLCFAs), increased production of plasmalogens, and improvement in mitochondrial function. ATAD1 was effective in fibroblast and HEK cell models of PEX1 and PEX6 ZSDs, as well as *in vivo* in a whole animal zebrafish ZSD model.

Our findings extend previous observations of ATAD1 efficacy in a PEX3 cell model^7^, suggesting that ATAD1 overexpression may have broad therapeutic applicability for ZSDs. Importantly, our work suggests that the role of ATAD1 extends beyond correcting mitochondrial morphology, as previously described, and involves restoring peroxisome-related metabolic functions^7,22,23^. By reducing VLCFA accumulation and restoring plasmalogen synthesis, ATAD1 normalizes two of the most clinically relevant biochemical abnormalities in ZSD, both of which contribute to severe neurological and systemic phenotypes. The fact that these improvements were observed across 3 distinct PEX mutations indicates that ATAD1 targets a (or several) convergent pathological mechanism. This is hypothesized to include the mislocalization of peroxins to mitochondria and the consequent disruption of mitochondrial integrity.

The loss of PEX1 or PEX6 severely impairs mitochondrial bioenergetics, and reintroducing these peroxins does not fully restore respiratory functions. This suggests that ongoing peroxisomal dysfunction leads to sustained mitochondrial damage, consistent with reports of disrupted lipid metabolism and redox imbalance in Zellweger spectrum disorders^2,24^. Such irreversible dysfunction may result from accumulated oxidative stress or defective protein import.

In contrast, ATAD1 overexpression significantly improved mitochondrial respiration in both PEX1^-/-^ and PEX6^-/-^ cells. ATAD1 has a role in removing mislocalized or damaged proteins from mitochondria. This may account for its ability to restore oxidative phosphorylation despite ongoing peroxisomal defects. Additionally, ATAD1 increased proton leak in PEX1 mutants, indicating mild uncoupling that helps protect against ROS^25^, and raised non-mitochondrial oxygen consumption, implying activation of compensatory redox pathways.

Overall, these results suggest a role for ATAD1 as a regulator of mitochondrial quality control, capable of restoring function even when peroxisomal recovery is incomplete. Targeting such proteostatic mechanisms may provide a therapeutic strategy for diseases involving organelle crosstalk and secondary mitochondrial dysfunction.

Our zebrafish Pex1 and Pex6 models provide further innovative tools to study ZSDs, including the molecular communication between peroxisomes and mitochondria. Both genes encode components of the AAA-ATPase complex, which is essential for importing proteins into the peroxisomal matrix^17^. Mutations in PEX1 or PEX6 are the most common causes of ZSDs^26^. Functionally, PEX1 and PEX6 form a hetero-hexameric complex that collaborates to dissociate PEX5 from the peroxisomal membrane, a critical step in recycling the import receptor and ensuring correct trafficking of matrix proteins. Because of this vital interaction, developing zebrafish lines lacking both genes offers the chance to explore whether simultaneous disruption of PEX1 and PEX6 leads to additive, synergistic, or redundant effects compared to losing just one gene.

Interestingly, despite the central role of the PEX1–PEX6 complex in peroxisomal biogenesis, our zebrafish locomotor assays at 6 dpf showed no significant behavioral impairment in single or double mutants under standard testing conditions. This result suggests that early zebrafish development may be protected against peroxisomal dysfunction, either through maternal contributions^27^, compensatory pathways^28^, or relatively low metabolic demands at this stage^29^. Phenotypic differences may become more pronounced under metabolic stress or at later developmental stages, when energy and lipid balance become increasingly crucial and the maternal yolk contribution is no longer available. Therefore, the double mutant line is advantageous because it allows investigations during embryo and larval stages, when zebrafish are most tractable, of genetic interactions during stress conditions or disease-related situations and potentially revealing deficits that are hidden in single mutants.

The Seahorse assays in Pex1-deficient zebrafish indicate a key vulnerability in mitochondrial adaptability. While basal and ATP-linked respiration were unaffected, Pex1 mutants exhibited a pronounced reduction in maximal respiratory capacity following FCCP treatment, which was restored by ATAD1 transient overexpression. This selective impairment in mutants suggests that mitochondrial plasticity, rather than baseline function, is compromised. Such a deficit aligns with the hypothesis that peroxisomal dysfunction does not immediately collapse mitochondrial metabolism but limits the ability of mitochondria to respond to energetic challenges^15,30,31^. This is consistent with reports that peroxisomes support mitochondrial performance by supplying lipids, buffering ROS, and facilitating redox balance^31,32^.

Although gene replacement therapy is an appealing option for ZSDs, a major limitation has been the size of the PEX genes, which exceed the capacity of adeno-associated virus (AAV)-mediated delivery^10,31^. Additionally, with at least 14 known PEX genes, each form of ZSD would need a lengthy and costly development process. Our studies with ATAD1 suggest a promising alternative. We have shown that the mistargeting of peroxins (PEX) contributes to the mitochondrial phenotype of PEX3^-/-^ ZSD, and that the extraction and degradation of mislocalized PEX, as well as other mislocalized peroxisomal proteins, is facilitated by ATAD1^7^. We now demonstrate that overexpressing ATAD1 is effective in PEX1 and PEX6 patient cell lines, as well as in vivo in zebrafish. Furthermore, the use of ATAD1 offers two significant and attractive features for gene therapy in ZSDs. First, its small size (∼1kb) enables efficient packaging in AAVs, whereas PEX1 and other PEX genes are larger than 4kb and, combined with necessary regulatory elements, exceed the AAV packaging capacity (∼4.8kb). Second, ATAD1 could potentially treat multiple different ZSDs. Finally, identifying ATAD1 as a shared downstream target for ZSDs provides a new conceptual framework, where a single pathway could potentially address various gene defects.

## ACKNOWLEDGEMENTS

Ian MacDonald and Matthew Benson generously shared the PEX6 cell line.

## METHODS

### Mammalian cell lines and culture

Human PEX1-deficient fibroblasts (GM16512) were obtained from the Coriell Cell Repository (Camden, NJ, USA). This line carries compound heterozygous mutations in PEX1, namely c.2097dupT (p.I700fs) and c.2528G>A (p.G843D). PEX6-deficient HEK293 cells, harboring a homozygous single-base deletion in exon 1 of PEX6 (c.544del) that causes a frameshift predicted to abolish protein function, were kindly provided by Benson et al^12^. Cells were cultured in Dulbecco’s Modified Eagle Medium (DMEM, high glucose, 4.5 g/L; Gibco) containing L-glutamine, supplemented with 10% heat-inactivated fetal bovine serum (FBS) and 1 mM sodium pyruvate. No antibiotics were used to avoid potential metabolic interference. Cultures were maintained at 37 °C in a humidified atmosphere of 5% CO₂ and regularly monitored for morphology and confluency.

### Retroviral production and transduction

HEK293T cells were maintained in DMEM supplemented with 10% FBS at 37 °C and 5% CO₂ and plated at 33–50% confluency in 100 mm dishes. Cells were transfected with plasmids encoding PEX6, PEX1, or ATAD1 together with pUMVC-GagPol and pCMV-VSV-G using Lipofectamine 3000 and P3000 reagent in Opti-MEM. After 24 h, viral supernatants were collected, clarified by centrifugation, filtered through 0.45 µm membranes, and used immediately for infection. Target PEX6-deficient HEK293 cells or PEX1-deficient fibroblasts were transduced with viral supernatant supplemented with 10 µg/mL polybrene in a 1:1 mixture with fresh DMEM + 10% FBS. Forty-eight hours post-infection, cells were selected with puromycin (0.5 µg/mL for PEX6 and PEX1 rescue) or Zeocin (125 µg/mL for ATAD1 rescue) for 4–7 days. All viral work was performed under biosafety cabinet conditions with appropriate personal protective equipment, and all materials were disinfected with ≥10% bleach after use.

### Lipidomic assays

We employed a multi-platform lipidomics workflow that combines both untargeted and targeted mass spectrometry to profile cellular lipids thoroughly. Lipids were extracted from cultured cells using a Matyash liquid–liquid extraction method, with the addition of class-specific, isotopically labeled internal standards for normalization across lipid classes. The organic phase was initially analyzed by quadrupole time-of-flight (QTOF) mass spectrometry for untargeted profiling of intact lipids and their fragments. Accurate mass measurements and MS/MS spectra supported the putative identification of lipid species, including fatty acids, cardiolipins, and triacylglycerols. These findings, along with prior experimental knowledge, guided the development of a targeted lipidomics panel. Targeted quantification was performed on two triple quadrupole (QQQ) instruments in multiple reaction monitoring (MRM) mode, utilizing complementary chromatographic separations to measure phospholipids, cholesteryl esters, sphingolipids, carnitines, and diglycerides. Quality control (QC) measures included pooled QC samples, replicate injections, and solvent blanks to evaluate reproducibility and carryover. Data from both platforms were integrated, before conducting statistical and enrichment analyses with MetaboAnalyst and LION/web. To standardize lipid names and ensure consistent structural annotation, we created a uniform nomenclature table using GOSLIN (Grammar of Succinct Lipid Nomenclature), enabling accurate downstream analysis and database integration.

### Peroxisome immunofluorescence staining and quantification

For peroxisome staining, HEK cells were cultured on 4-well chamber slides and fixed with freshly prepared 4% paraformaldehyde (PFA) for 30 minutes at room temperature (RT). After fixation, wells were washed three times with Dulbecco’s Phosphate-Buffered Saline (DPBS), and cells were blocked for 30 minutes at RT using a blocking solution made of 1% bovine serum albumin (BSA) and 0.1% Tween-20 in DPBS.

Cells were incubated with rabbit anti-PMP70 primary antibody (Abcam, ab109448) diluted 1:200 in DPBS for at least 45 minutes at room temperature with gentle rocking. After primary incubation, slides were washed three times with DPBS (5 minutes each) and incubated with secondary antibody (goat anti-rabbit Alexa Fluor 488 or Alexa Fluor 555) diluted 1:400 in DPBS for 30 minutes at room temperature in the dark. Slides were then washed three more times with DPBS (5 minutes each). After the final wash, 500 µL of Fluoromount-G was added to each well for mounting.

PMP70⁺ area (µm²) and PMP70⁺ area as a percentage of total cell area were measured using QuPath software (https://qupath.github.io, version 0.2.3). Images for QuPath analysis were captured with the Nikon Spinning Disk Confocal microscope.

### Mammalian cell lines Seahorse Assay

Mitochondrial oxygen consumption rate (OCR) was measured using the Seahorse XF Pro Analyzer (Agilent). Cells were seeded at a density of 10,000 cells per well in XF96 cell culture microplates (Agilent, Cat#101085) and maintained in XF-formulated Dulbecco’s Modified Eagle Medium (DMEM). Before the assay, plates were pre-warmed, and the analyzer temperature was set to 37 °C. OCR was recorded over cycles consisting of 3 minutes of mixing followed by 3 minutes of measurement. Basal respiration was measured over 3 cycles prior to drug injection.

Sequential compound injections were performed using an injection volume of 20 µL per port to achieve final well concentrations of 3 µM oligomycin (port A), 1.5 µM FCCP (port B), and 3 µM each of rotenone and antimycin A (port C). Each injection step was followed by 3 measurement cycles. Oligomycin was used to inhibit ATP synthase and determine ATP-linked respiration and proton leak. FCCP was then added to uncouple oxidative phosphorylation and assess maximal respiratory capacity. Finally, rotenone and antimycin A were injected to inhibit complexes I and III, enabling the quantification of non-mitochondrial respiration.

### Generation of the zebrafish Pex1 mutant line

Using CRISPR/Cas9, we targeted *Pex1* (NM_001170835.1). A single guide RNA (sgRNA) was synthesized using a cloning-free method based on a protocol described previously. The guide portion of the sgRNA, targeting a portion of exon 5 (AGATGGCTGACAGACCTAGG), was designed using CHOPCHOP. The sgRNAs were prepared as previously described [30], resulting in a fragment consisting of the T7 promoter, the gene-specific sequence, and the guide core sequence. The sgRNAs were synthesized by in vitro transcription using the HiScribe T7 High Yield RNA Synthesis Kit. RNAs were purified using the RNA Clean & Concentrator-5 kit (Zymo Research), eluted with 20 μL of water, and diluted to working concentrations of ∼ 400 ng/μL.

### Founder screening and identification of heterozygous adult fish

sgRNA was injected, and the fish were genotyped for potential mutations at the target site. One strain with an allele containing a 44 bp deletion, resulting in a premature stop codon, was identified via Sanger sequencing and HRM analysis and was selected for further experiments. The identified F1 founders were crossed with wild-type zebrafish (TL strain), and their adult offspring (F2) were genotyped. Heterozygous F2 fish from the mutant line were crossed, and the offspring were observed using bright-field and fluorescence microscopy. Embryos displaying phenotypes were euthanized with an overdose of MS-222 (300mg/L) and genotyped by HRMA (forward primer: 5’-GCCATCCTGAAAGACTGTATCC-3’; reverse primer: 5’-CTCTTGCTCTTGAAGGTTCCAT-3’).

### Zebrafish behavioral assays

Larvae were separated into 24-well plates and then incubated in a 28°C incubator for 20 minutes. Larval zebrafish locomotor activity was analyzed for 30 minutes using the DanioVision system and Noldus Ethovision XT 13 program (Noldus, Wageningen, the Netherlands) for larval behavioral assay. Larvae were randomly analyzed on the same plate, and HRMA was used to determine their genotype after the experiment was completed. All behaviors were analyzed between 9:00 a.m. and 5:00 p.m.

### Zebrafish Seahorse assay

At 1dpf, embryos were manually dechorionated using fine forceps. They were then maintained at 28.5 °C in E3 medium until 2 dpf, at which point they were anesthetized with 125 µg/mL tricaine (MS-222). For respirometry, each embryo was transferred individually into a well of a Seahorse XF96 cell culture microplate (Agilent, Cat#101085), with each well containing 170 µL of E3 medium. Transfers were performed using a 100 µL pipette tip to reduce mechanical stress. The plates were pre-warmed, and the Seahorse XF Pro Analyzer temperature was set to 28.5 °C. Basal respiration was measured over 12 cycles, each consisting of 2 minutes of mixing followed by 2 minutes of measurement. Using an injection volume of 2.28 µL per port, the final concentrations in the well were: 12 µM oligomycin (port A), 2.5 µM FCCP (port B), and 1 µM rotenone plus 1 µM antimycin A (port C). After baseline cycles, oligomycin was injected to inhibit ATP synthase, and ATP-linked respiration and proton leak were measured over 20 cycles. FCCP was then added to uncouple oxidative phosphorylation and assess maximal respiratory capacity over 8 cycles. Finally, rotenone and antimycin A were injected sequentially to inhibit complexes I and III, allowing determination of non-mitochondrial respiration.

### ATAD1-2A-eGFP for Capped mRNA Synthesis

To generate capped mRNA encoding ATAD1 fused to eGFP via a 2A self-cleaving peptide, two overlapping DNA fragments were amplified by PCR and subsequently fused by overlap extension. The resulting full-length construct served as the template for in vitro transcription. Four primers were used to amplify and assemble the fusion construct. For the ATAD1 fragment, the forward primer SP6-Kozak-ATAD1_1F (5′-ATTTAGGTGACACTATAGCCACCATGGTACATGCTGAAGCCTT-3′) and the reverse primer 2A-ATAD1_2R (5′-AAGTTCGTGGCTCCGGATCCTAAACAAACATGTGTTAAAACATTCTG-3′) were used.

To amplify the 2A-eGFP-pA fragment, the forward primer ATAD1-2A_3F (5′-CAGAATGTTTTAACACATGTTTGTTTAGGATCCGGAGCCACGAACTT-3′) and the reverse primer pA_4R (5′-AAAAAACCTCCCACACCTCC-3′) were employed. PCR products were resolved on a 1% TAE agarose gel, and the expected bands were excised and purified using the Zymo Gel DNA Recovery Kit. To generate the full-length fusion construct, overlap extension PCR was performed using the purified ATAD1 and 2A-eGFP-pA fragments as templates, with SP6-Kozak-ATAD1_1F and pA_4R as external primers. The resulting product was gel-purified and quantified. The purified fusion construct was used as a template for in vitro transcription using the mMESSAGE mMACHINE™ SP6 Transcription Kit (Thermo Fisher Scientific), according to the manufacturer’s instructions. This reaction produced capped mRNA, which was subsequently purified and stored at −80°C until it was injected into *Pex1* zebrafish embryos.

### Zebrafish and mammalian cell lines quantitative real-time PCR

For zebrafish experiments, total RNA was extracted from five pooled larval fish using the PureLink RNAmini Kit (Invitrogen). For tissue culture experiments, total RNA was extracted from cell pellets of different biological replicates containing 1 × 10⁶ cells. cDNA was synthesized using SuperScript™ III reverse transcriptase (Invitrogen). Real-time PCR was conducted on a QuantStudio 12K Flex Real-Time PCR System (Thermo Fisher Scientific) with the PowerUp SYBR Green master mix (Thermo Fisher Scientific) according to the manufacturer’s instructions. mRNA levels were normalized to the average of β-actin in zebrafish and GAPDH in cells, serving as internal controls. The qPCR analysis was performed using the 2 − ΔΔCT method.

### AlphaFold Modeling

Predicted structural models for human PEX1 (UniProt: O43933), human PEX6 (UniProt: Q13608), zebrafish Pex1 (UniProt: D2CK11), zebrafish Pex6 (UniProt: F1QMB0), and all derived mutants were acquired from AlphaFold (https://alphafoldserver.com/) and visualized using The PyMOL Molecular Graphics System, Version 2.5, Schrödinger, LLC.

